# 4-Nitrobenzoate inhibits 4-hydroxybenzoate polyprenyltransferase in malaria parasites and enhances atovaquone efficacy

**DOI:** 10.1101/2025.06.10.658244

**Authors:** Ignasi Bofill Verdaguer, Matheus Felipe Santos, Maurício Mazzine Filho, Gabriela Oliveira Castro, Agustín Hernandez, Manoel Aparecido Peres, Alejandro Miguel Katzin, Marcell Crispim

## Abstract

Ubiquinone (UQ) is a critical component of the electron transport chain in *Plasmodium falciparum*, the etiological agent of human malaria. The first step in UQ biosynthesis is the condensation of 4-hydroxybenzoate (4-HB) and an isoprenic chain by the enzyme 4-hydroxybenzoate polyprenyltransferase (4-HPT; COQ2 gene). Atovaquone (AV), an antimalarial drug, competes with ubiquinol (UQH_2_) for binding to the mitochondrial bc1 complex, preventing the redox recycling of UQ. In clinical practice, AV is combined with proguanil, a dehydrofolate reductase inhibitor, in a single formulation. However, parasitic resistance to this combination has been demonstrated, indicating the need for new pharmacological combinations to potentiate AV. Previously, 4-nitrobenzoate (4-NB) demonstrated the ability to inhibit UQ biosynthesis in *P. falciparum* parasites as well as potentiate AV efficacy *in vitro*. However, both its pharmacodynamics and whether this potentiation could be useful *in vivo* remained obscure. Here we show that 4-NB enhances AV antiplasmodial efficacy to kill parasites, increases its selectivity compared with animal cells, and preserves proguanil efficacy. 4-NB specifically inhibited the 4-HPT enzymatic activity in mutant strains of *Saccharomyces cerevisiae* complemented with *Pf*COQ2. Finally, 4-NB also improved AV antimalarial efficacy in mice infected with *Plasmodium berghei* parasites. Finally, work with various 4-HB analogs delineated the chemical requirements to potentiate AV activity. These findings clarify the importance of UQ biosynthesis for malaria parasites and suggest that *Pf*COQ2 could be a therapeutic target to enhance the efficacy of AV.

## 1. Introduction

Malaria is one of the most widespread parasitic diseases in tropical and subtropical regions, affecting millions of people every year (WHO, 2023). Most deaths occur in Africa due to *Plasmodium falciparum* infections. Several drug resistances have already been reported, and new antimalarial therapies are required (Färnert et al., 2003; Farooq & Mahajan, 2004). Atovaquone (AV) is one of the treatments for malaria. This compound competes with ubiquinol (UQH_2_) for binding to the cytochrome bc1 complex, thereby preventing ubiquinone redox recycling (UQ) (Radloff et al., 1996; McKeage et al., 2003; Verdaguer et al., 2021). As a consequence, the activity of dihydroorotate dehydrogenase (DHODH), involved in pyrimidine biosynthesis, is halted, leading to an inviable condition for the parasite. ATP production via oxidative phosphorylation does not seem to be essential in intraerythrocytic stages (Painter et al., 2007). In the clinic, AV is synergistically combined with proguanil in a single formulation, Malarone®. Albeit a known inhibitor of folate biosynthesis, proguanil has been previously demonstrated to help AV to collapse the mitochondrial membrane potential, possibly by lowering the levels of ubiquinone; however, the precise mechanism by which it happens remains unknown (Srivastava & Vaidya, 1999). In any case, parasitic resistance to AV plus proguanil has also been found, and new pharmacological combinations to potentiate AV are required (Vaidya et al., 2000; Färnert et al., 2003).

*P. falciparum* biosynthesizes UQ-8 and UQ-9, while humans mostly produce UQ-10 (Ellis, 1994; de Macedo et al., 2002; Verdaguer et al., 2019). UQ biosynthesis occurs in the mitochondria and starts with the condensation of 4-hydroxybenzoate (4-HB) and an isoprenic chain by the transmembrane enzyme 4-hydroxybenzoate polyprenyltransferase (4-HPT), encoded by the *COQ2* gene (Figure 1). The resulting compound, 3-polyprenyl-4-hydroxybenzoate, is enzymatically modified by hydroxylations, decarboxylations, and three S-adenosyl-L-methionine (SAM)-mediated methylations, leading to UQ formation. In malaria parasites, 4-HB for UQ aromatic head group biosynthesis is allegedly provided by the shikimate pathway and, possibly, exogenous incorporation (Valenciano et al., 2019). On the other hand, the Methyl Erythritol Phosphate (MEP) pathway seems to be the source of isoprenoids for UQ lateral chain formation (Verdaguer et al., 2019). The MEP pathway is located in a non-photosynthetic plastid called the apicoplast, which leads to the production of two 5-carbon molecules, isopentenyl pyrophosphate (IPP) and its isomer dimethylallyl pyrophosphate (DMAPP). Both substances can be subsequently condensed for the formation of longer polyprenyl moieties, including geranyl pyrophosphate (GPP, 10 carbons); farnesyl pyrophosphate (FPP, 15 carbons); geranylgeranyl pyrophosphate (GGPP, 20 carbons); and those directly employed for UQ biosynthesis, octaprenyl-PP and nonaprenyl-PP (Verdaguer et al., 2019).

**Figure 1.**
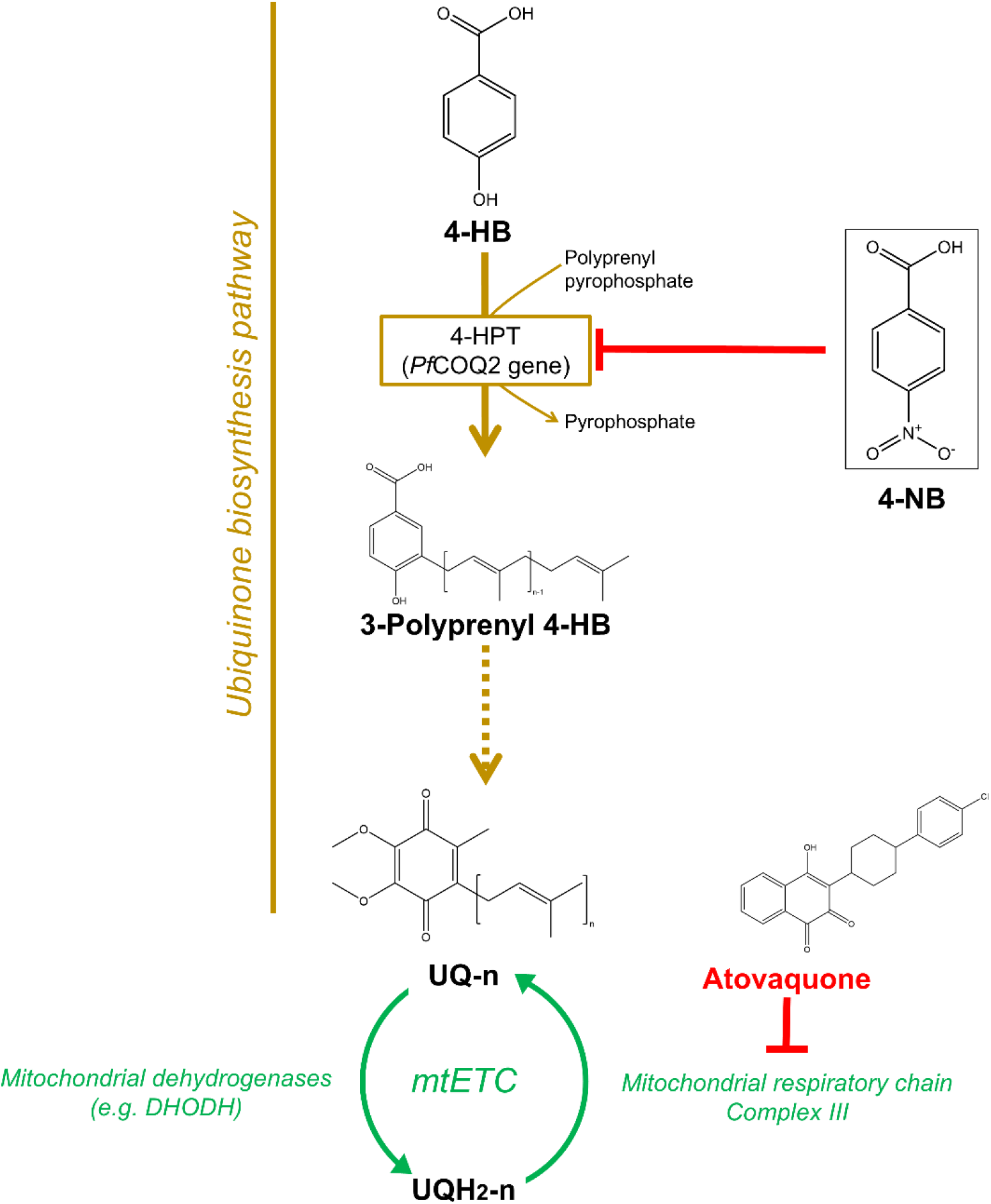
Ubiquinone biosynthesis pathway. Schematic representation of the UQ biosynthetic pathway (brown), showing the chemical structures of precursors, intermediates, and UQ itself. The enzyme 4-HPT (EC 2.5.1.39), the hypothetical target of 4-NB in malaria parasites, is indicated in *italic*. Discontinuous lines represent multiple steps; red arrows indicate drug inhibition. The chemical structures of 4-HB and 4-NB are boxed. The figure also shows UQ redox recycling in the mitochondrial electron transport chain (mtETC, green), and the chemical structure and molecular target of atovaquone.

Previous studies in our group revealed that [^3^H] GGPP incorporation into UQ in malaria parasites can be inhibited by 4-nitrobenzoate (4-NB). Interestingly, although 4-NB demonstrates poor antiplasmodial effects on its own, it has a significant ability to potentiate the efficacy of AV *in vitro*. Presumably, a depletion of the UQ pool would facilitate the AV interaction with mitochondrial complex III and thus, diminish UQ-redox regeneration required for DHODH activity. In any case, this potentiation effect is reduced when 4-HB is added to the medium indicating that the target of 4-NB is related to UQ biosynthesis (Verdaguer et al., 2021). In parallel to this, we also showed that the gene PF3D7_0607500 (*PfCOQ2*) can complement the *COQ2* gene of *S. cerevisiae*, demonstrating that *PfCOQ2* encodes for a functional 4-HPT enzyme (Zafra et al., 2023). Consistent with these findings, previous studies reveled that the product of *PfCOQ2* is located in the parasitic mitochondria (Van Dooren et al., 2006). Despite all these findings, the enzymatic target of 4-NB remained unknown and its utility for *in vivo* use unexplored. In this study, we found that the molecular target of 4-NB seems to be the 4-HPT enzyme and that 4-NB not only enhanced the *in vitro* antiplasmodial efficacy of AV but it also increased its selectivity compared to animal cells while still preserving the AV-proguanil synergistic interaction. Additionally, 4-NB boosted AV efficacy in treating murine malaria.

## 2. Methods

### 2.1 Reagents and stock solutions

[ring-^14^C (U)] 4-HB (50-60 mCi/mmol) was purchased from American Radiolabeled Chemicals (Saint Louis, Missouri, USA). RPMI-1640, medium and Albumax I (0.5%) and heat-inactivated Bovine Fetal Serum (BFS) were obtained from Thermo Fisher Scientific (Waltham, Massachusetts, USA). All HPLC grade solvents, AV, proguanil hydrochloride, and other reagents not cited here were purchased from Sigma (St. Louis, Missouri, USA). SYBR Green I® nucleic acid gel stain from Life Technologies® (Eugene, OR, USA), and UQ-8,9 standards were purchased from Avanti Polar Lipids (Alabaster, Alabama, USA). Sterile stock solutions for *in vitro* use were prepared: 25 mM AV in dimethyl sulfoxide, filter-sterilized 800 mM proguanil hydrochloride in water, 4-HB and all its analogues were dissolved at 100 mM in ethanol.

### *2*.*2 Plasmodium falciparum* asexual stages culture

The *P. falciparum* 3D7 strain was cultured in 75 cm^2^ flasks using RPMI-1640 medium complemented with Albumax I (0.5%) and a gaseous mixture of 5% CO_2_, 5% O_2_, and 90% N_2_ obtained from Air Products Brasil LTDA (São Paulo, SP, Brazil) following the Trager and Jensen methodology (Trager and Jensen 1976; Crispim et al., 2022). Culture synchronization was performed using 5% (w/v) D-sorbitol solution as previously described (Lambros & Vanderberg, 1979), and parasitic stages and parasitemia were monitored by Giemsa-stained smears microscopy. To avoid culture contamination, PCR tests for mycoplasma were regularly carried out (Rowe et al., 1998).

### 2.3 Culture and complementation of *COQ2* gene in yeasts

The culture and introduction of *COQ2* genes in yeast were performed as previously described (Zafra et al., 2023). Cells were cultured in YPD medium (2% glucose, 2% peptone, 1% yeast extract), YPGly medium (2% glycerol, 2% peptone, 1% yeast extract), or Synthetic Defined medium (SD) without uracil, and with 2% glucose or 3% glycerol as a carbon source. The *ΔCOQ2 S. cerevisiae* strain (YNR041C, MATa *ade2-1 his3-1,15 leu2-3,1 12 trp1-1 ura3-1 COQ2:HIS3*) and its parental W303-1a were generously provided by Dr. M. H. de Barros (Dept. Microbiology, University of São Paulo). The *ΔCOQ2* yeast strain was complemented with either its own gene (*ScCOQ2*; YNR041C; positive control) or PF3D7_0607500 gene (referred to as Pf*COQ2* hereafter) optimized for its expression in yeast by Genscript. In both cases, genes were cloned into a p416-GPD yeast expression vector and introduced into yeast using the lithium acetate method (Gietz & Woods, 2002). The phenotype was evaluated as ability to use non-fermentable (glycerol) or fermentable (glucose) carbon sources (Zafra et al., 2023).

### 2.4 Yeast growth tests

Drop growth tests were conducted on agar plates with SD medium without uracil, using 2% glucose or glycerol as the carbon source, and with or without the addition of 1 mM 4-HB, 1 mM 4-NB, or a combination of 1 mM 4-HB and 1 mM 4-NB. Both 4-HB and 4-NB were added from a 100-fold concentrated stock solution prepared in ethanol. For drop tests, cells from each strain were grown to early stationary phase in SC medium without uracil and with 2% glucose. The culture absorbance was then adjusted to 0.4 (approximately 4 × 10^6 cells ml−1) using sterile water, and several 10-fold serial dilutions in water were prepared from the initial culture. Aliquots of 2.5 μl from each dilution were placed sequentially onto appropriate agar Petri dishes. The inoculated plates were then incubated at 30°C for 48-72 hours before being photographed.

### 2.5 4-hydroxybenzoate polyprenyltransferase activity

The *in vitro* function of 4-HPT was characterized using crude membrane extracts from the yeast *ΔCOQ2* strain transformed with p416-Pf*COQ2*. For this, yeasts were cultured to early stationary phase in SD plus glucose medium, and then cells were disrupted by glass beads (0.5 mm diameter) (Dunn & Wobbe, 1993). Unbroken cells were removed by centrifugation at 900 x *g* for 1 min, and protein was adjusted to 50 mg/mL with 100 mM Tris/HCl pH 7.5. Commercially available FPP was used as isoprenic donor, as most 4-HPT enzymes studied show broad substrate specificity for the isoprenic chain length (Melzer & Heide, 1994; Sies & Packer, 2004). The reaction was performed in 1.5 mL microtubes by incubating 4 mg of yeast protein with 10 mM MgCl_2_, 50 µM FPP, and 10 µM [ring-^14^C (U)] 4-HB. The volume was adjusted to 100 µL with 100 mM Tris/HCl pH 7.5, and the reaction was initiated by adding the yeast extract. In some assays, 4-HB analogues were also added to the reaction, prenyl-PP was omitted, or boiled yeast extracts were employed as controls. All the substances diluted in ethanol were dried under a vacuum to avoid introducing solvents into the reaction (this includes drugs and [ring-^14^C (U)] 4-HB). After 1 h of incubation at 37°C, the reaction was stopped by adding 200 µL of ethyl acetate. The mixture was vortexed, centrifuged at 12,000 *x g* for 10 min, and the organic phase was dried under vacuum. The residue was suspended in 10 µL of ethyl acetate and chromatographed on silica 60 plates (20×20 cm, Merck). Plates were developed with acetone: petroleum ether (7:3, by volume) (Pfaff et al., 2014). Authentic standards of 4-HB and pABA were run on the same plate. Standards were visualized with iodine vapor. Finally, the plates were treated with EN3HANCE (Perkin Elmer) and exposed to autoradiography for one week at −70°C. Products of the reaction were identified by their Rf and the controls previously described (Pfaff et al., 2014).

### 2.6 Parasitic growth monitoring

The antimalarial effects of the compounds, separately or in combination, on parasitic growth were monitored relative to an untreated control using previously published methods (Desjardins et al., 1979; Smikstein et al., 2004). The experiments were done in 96-well plates starting at the ring stage (2% parasitemia, 2% hematocrit). Several concentrations of each compound were prepared by serial dilution in RPMI complete medium. Parasite growth was monitored after 48 hours by SYBR Green I® DNA staining as described elsewhere (Smilkstain et al., 2004). Briefly, 100 μL of culture was incubated in a 96-well cell plate in the darkness and at room temperature after adding 100 μL of Syber Green I® 1/5,000 (v/v) in lysis buffer [20 mM Tris, pH 7.5; 5 mM EDTA; 0.008% saponin (w/v); 0.08% Triton X-100 (v/v)]. Fluorescence was measured in a POLARstar Omega fluorometer® (BMG Labtech®, Ortenberg, Germany) with excitation and emission bands centred at wavelengths of 485 and 520 nm, respectively.

### 2.7 Assessment of antiplasmodial drug effects and drug interaction studies

The concentration of each compound required to decrease parasitic growth by 50% (IC_50_) was determined at 48 hours. Inhibition of parasite growth was analyzed in relation to the logarithm of the concentration using a nonlinear regression (dose-response slope / variable sigmoid equation) with GraphPad Prism® software. All experiments monitoring parasitic growth were performed at least three times with four technical replicates for each one. For the study of AV interaction with compounds with poor antimalarial activity it was employed a potentiation assay, as described elsewhere (Verdaguer et al., 2021). In this case, the AV IC_50_ value was calculated at a fixed sublethal concentration of different compounds. The single-drug fractional inhibitory concentration value (FIC value) was calculated as the IC_50_ value of AV in a combined solution divided by the IC_50_ value of AV alone. AV FIC < 0.5 was considered indicative of a drug potentiation phenomenon (Verdaguer et al., 2021).

### 2.8 *In vivo* experiments

Three female Balb/c mice, aged 4-6 weeks, were used for each group, and experiments were performed three times. Mice were infected by intraperitoneal injection with 1 × 10^7^ infected erythrocytes from a donor mouse. The animals were kept in standard conditions with a 12-hour light/dark cycle, in cages with autoclaved pine wood and free access to water and food. On the fourth day post-infection, administration of 4-NB or AV at subtherapeutic doses was initiated. 4-NB was administered at 88 mg/kg from a 5.3 g/L PBS sterile stock solution. The 4-NB stock was prepared by dissolving the compound in PBS and adjusting the pH to 7.4 with sodium hydroxide until complete solubilization. Then, DMSO was added to a final concentration of 0.1%. The chosen dose of 4-NB corresponds to one-tenth of the lethal dose 50 (Mandel & Metais, 1948). AV was administered at 0.1 mg/kg from a 20 µg/mL sterile stock solution (Nuralitha et al., 2017). The AV stock was prepared by dissolving 10 mg AV in 1 mL DMSO and then diluting it 1000-fold in PBS for IP administration. For combinatory treatments, AV dissolved in DMSO was diluted in the previously described 4-NB stock solution. In all cases, the drugs were administered once daily, every 24 hours, over a period of 5 days. Parasitemia was monitored through tail blood smears every two days, and mice were euthanized when it exceeded 40%. To compare the means of variables, the unpaired student’s t-test was employed. In addition, survival curves were plotted using the Kaplan-Meier method, and statistical differences were evaluated using the log-rank test. The analyses were performed using the GraphPad PRISM® 5.3 program.

### 2.8 Animal cells culture and growth monitoring

The kidney epithelial cells from *Macaca mulatta* (LLC-MK2) were grown routinely in 75 flasks in RPMI medium supplemented with 10% FBS and 10 mg/L gentamicin sulfate. The cultures were maintained in a humidified incubator with 5% CO_2_ at 37 °C. Cells were manipulated following the passage and trypsinization procedures as described elsewhere (Mosmann et al., 1983; Arodin et al., 2019). For experiments, confluent cultures were washed in Phosphate Buffered Saline (PBS), trypsinized, centrifuged at 300 x *g*, and suspended in culture media. The cells were cultured in 96-well plates at a density of 1.0 × 10^5^ cells/well. The next day, the cells were washed with PBS and subjected to pharmacological treatments. Ethanol controls were included to take into consideration any effects related to solvents, and its concentration was always ≥1%. After 48 hours, the cells were washed once in PBS, and each well was incubated at 37°C with 50 µL PBS containing 5 mg/mL MTT. After 4 hours, 50 µL of 20% SDS in PBS (wt/vol.) was added to each well. The next day, the absorbance at 595 nm wavelength corrected to 690 nm was monitored in a POLARstar Omega fluorometer® (BMG Labtech®, Ortenberg, Germany), and the results were analyzed by GraphPad Prism® software to determine the 50% cytotoxic concentration (CC_50_) (Mosmann et al., 1983). Statistical significance was determined by Student’s t-test, one-way ANOVA, or nonlinear regression (dose-response). All experiments were repeated at least three times, with four technical replicates each time, and PCR for mycoplasma and optic microscopy were used to avoid culture contamination (Nikfarjam et al., 2012). Selectivity Index (SI) was calculated as CC_50_ in cells / IC_50_ in parasites. As other authors have described, a selectivity index >10 indicates promising antibiotic potential (Kissin, 2013).

## 3. Results

### 3.1 4-Hydroxybenzoate analogues inhibit recombinant *PfCOQ2*

As the enzymatic target of 4-NB in *P. falciparum* remains unknown, we hypothesized that it could be the 4-HPT enzyme. To assess this, a set of yeast strains were created to study pharmacological inhibition of the malaria parasite 4-HPT enzyme. The W303-1a *S. cerevisiae* strain was transformed with the p416-GPD vector (referred to as plasmid control, PC), and the ΔCOQ2 strain was transformed with either the p416-GPD empty plasmid (referred to as the ΔCOQ2-0 strain), the p416-GPD-ScCOQ2 plasmid (referred to as the ScCOQ2 strain), or the p416-GPD-PfCOQ2 plasmid (referred to as the PfCOQ2 strain). The phenotype of these strains was assessed by their ability to use a non-fermentable carbon source (SD + glycerol) in the presence or absence of different combinations of 4-HB and 4-NB (Figure 2A; see another replicate in Supporting Information, Figure S1). As expected, the ΔCOQ2-0 strain was only able to grow on SD + glucose plates, while all strains expressing COQ2 genes were able to grow on both SD + glucose and SD + glycerol plates. The addition of 1 mM 4-NB negatively affected the growth of all strains in SD + glucose medium. In SD + glycerol, 4-NB completely inhibited the growth of the PfCOQ2 strain but not the ScCOQ2 strain. This may indicate that 4-NB has a mechanism of action related to respiratory metabolism and directly involves the 4-HPT enzyme of malaria parasites. The addition of 1 mM 4-HB to both media partially rescued the growth of the PfCOQ2 strain from the effects of 4-NB on SD + glycerol plates, indicating that 4-NB acts as an antimetabolite of 4-HB. 4-HB itself had no effect on yeast strains growth in either SD + glucose or SD + glycerol media.

**Figure 2.**
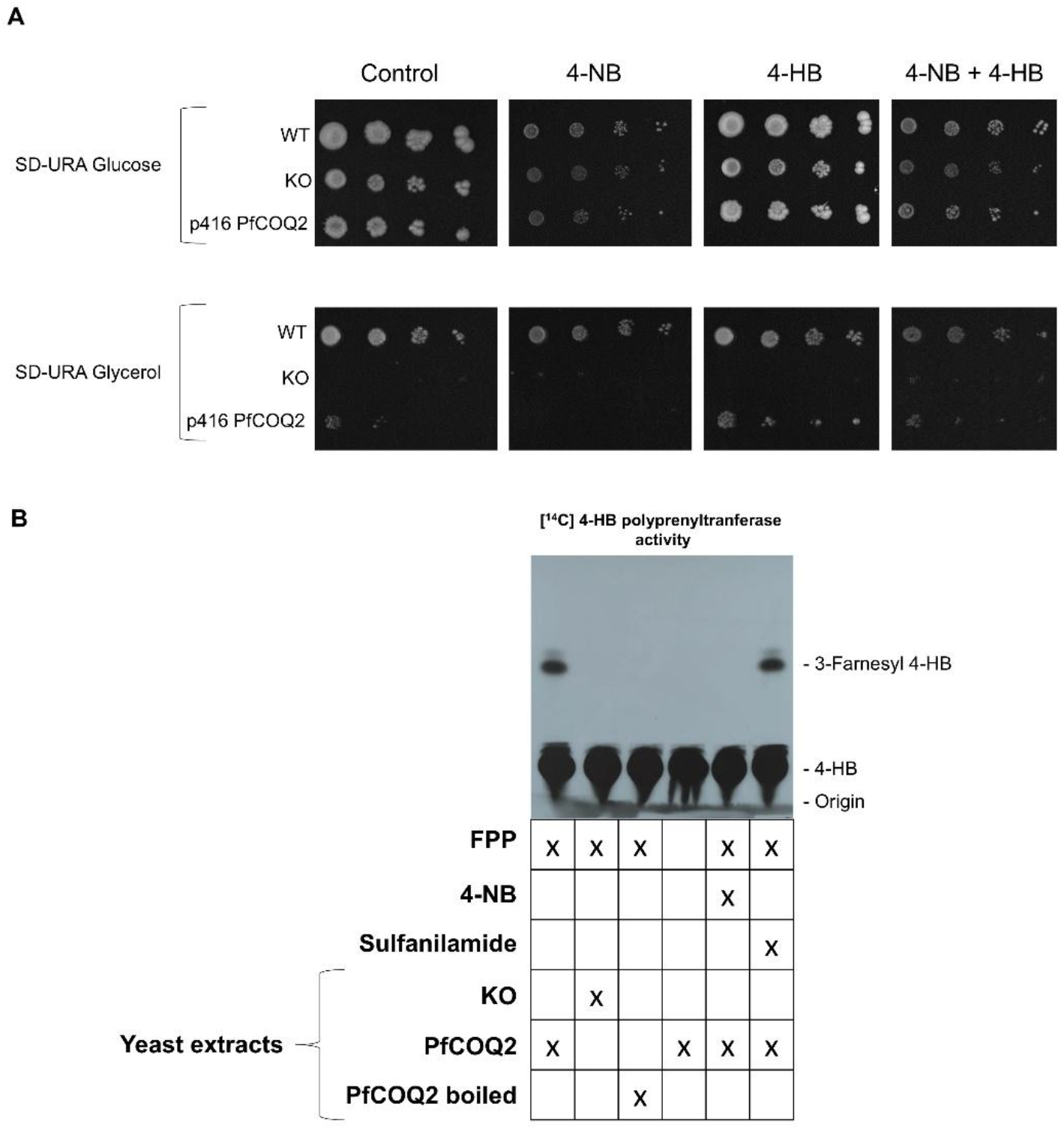
*PfCOQ2* complementation and enzymatic activity. (A) The figure shows the growth of yeast strains in SD + glucose / SD + glycerol media containing a concentration of 1 mM of different drugs, as indicated. This experiment was performed three times with similar results. (B) The figure shows the analysis of 4-HPT enzymatic activity in *Pf*COQ2 complemented yeasts under the presence of 0.5 mM of different drugs. The compounds added to the enzymatic reaction are indicated on the TLC autoradiography. The retention of different compounds is also indicated. This experiment was performed three times with similar results.

Considering the last set of results, we wanted to directly assess the enzymatic inhibition of Pf*COQ2* (Figure 2B; see another replicate in Supporting Information, Figure S1). When whole yeast extracts were incubated with [ring-^14^C (U)] 4-HB plus FPP, radiolabeled compounds were produced which were chromatographically compatible with 3-farnesyl 4-HB (Figure 2 B), as previously described (Pfaff et al. 2014). Other radiolabeled compounds were also observed, with one of them being identified as a contaminant present in the [ring-^14^C (U)] 4-HB reagent. The other compounds may be SAM-methylated derivatives of 3-polyprenyl 4-HB. This type of compound formation is commonly seen when assaying the product of *COQ2* activity in raw extracts (Sies & Packer, 2004). However, prenylated derivatives of [ring-^14^C (U)] 4-HB were not observed in Pf*COQ2* boiled extracts, extracts of ΔCOQ2-0 strain or assays with Pf*COQ2* strain without the addition of polyprenyl-PP, which indicates the enzymatic origin of these compounds as well as their prenylated nature. Furthermore, the addition of 0.5 mM 4-NB inhibited the prenylation of [ring-^14^C (U)] 4-HB. Similarly, the aromatic compound sulfanilamide, used as an aromatic-chemical control compound similar to 4-NB, did not inhibit this 4-HPT activity.

### 3.2 4-Nitrobenzoate enhances AV selectivity, potency and synergy with proguanil

Since both proguanil and 4-NB seem to inhibit UQ biosynthesis in malaria parasites, the effects of 4-NB on proguanil were investigated (Table 1). The IC_50_ values of AV and proguanil were calculated as a reference for further comparison. Then, the same assay was performed by supplementing the RPMI medium with 0.5 mM 4-NB, a concentration which has no effect on parasitic growth at 48 hours, as described elsewhere (Verdaguer et al., 2021). Results showed that 4-NB potentiates AV efficacy x 6.25-fold but not proguanil. These results demonstrate that 4-NB enhances the antiplasmodial efficacy of AV against parasites while maintaining the efficacy of proguanil. This suggests that the mechanisms by which proguanil and 4-NB potentiate AV are likely different.

**Table 1.** 4-NB effects on AV potency and selectively. The table shows the IC_50_/CC_50_ ± SD values for Proguanil and AV, as well as the SI values for AV, in the absence/presence of 0.5 mM 4-NB in *P. falciparum* parasites and LLC-MK2 cells. This experiment was performed three times.

|  | IC<sub>50</sub> proguanil (μM) in<br>malaria parasites | IC<sub>50</sub> atovaquone (nM) in<br>malaria parasites | CC<sub>50</sub> atovaquone (nM) in<br>LLC-MK2 cells | SI atovaquone |
| --- | --- | --- | --- | --- |
|  |  | IC<sub>50</sub> ± SD | CC<sub>50</sub> ± SD |  |
| Control | 18.76 ± 0.02 | 0.550 ± 0.012 | 5.105 ± 1.520 | 0.928 × 10<sup>4</sup> |
| 4-NB | 23.59 ± 3.39 | 0.088 ± 0.004 | 3.817 ± 2.012 (ns) | 4.34 × 10<sup>4</sup> |

Other authors demonstrated that AV also inhibited cytochrome bc1 and mtETC in mammalian cells (Fiorillo et al., 2016). Thus, it was decided to evaluate how a potential combination of AV with 4-HB analogues would affect the selectivity of the antimalarial. The presence of 0.5 mM concentration of 4-NB slightly reduced the CC_50_ value of AV in the LLC-MK2 cell line (Table 1). However, statistical analysis showed that none of the observed potentiating effects were significant. As previously seen, all 4-HB analogues triggered a reduction in the IC_50_ value of AV in the malaria parasite.

### 3.3 4-Nitrobenzoate enhances atovaquone efficacy *in vivo*

As observed, 4-NB enhanced the selectivity and potency of AV. Therefore, it was of interest to test this compound in mice, either alone or in combination with AV. For this purpose, animals were infected with *P. berghei* ANKA parasites and four days after were treated with suboptimal doses of AV (0.1 mg/kg, intraperitoneally) (Nuralitha et al., 2017), 4-NB (88 mg/kg, intraperitoneally) (Mandel & Metais, 1948) or a combination of both (0.1 mg/kg AV plus 88 mg/kg 4-NB, both administrated intraperitoneally). Compounds were administered every 24 h for 4 days, and a control group of untreated animals was also performed (Figure 3 and Supporting Information Figure S2). Compared to untreated animals, mice treated with AV alone exhibited a slight reduction in parasitemia levels and experienced a significantly extended survival. However, animals treated with AV in combination with 4-NB experienced a significantly more pronounced decrease in parasitemia, followed by a slower increase in parasitemia levels, and extended survival time compared to those animals that received only AV. Mice treated with 4-NB alone showed no reduction in parasitemia levels but a slightly reduced survival compared to untreated animals. These results indicate that 4-NB possesses no antimalarial activity by itself but enhances the efficacy of AV *in vivo*.

**Figure 3.**
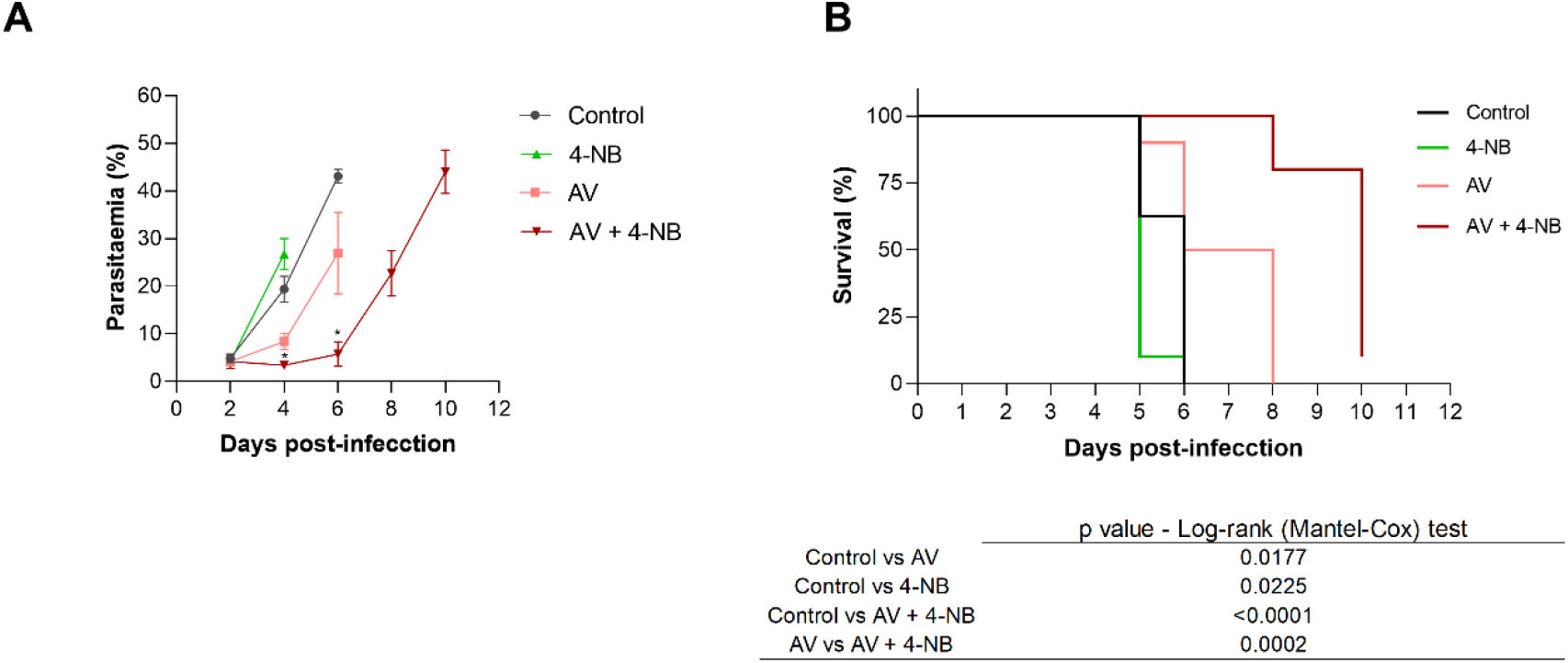
4-Nitrobenzoate improves the efficacy of atovaquone *in vivo*. (A) Parasitemia in *P. berghei* ANKA-infected mice treated with atovaquone (AV), 4-nitrobenzoate (4-NB), or their combination. All groups received the same amount of vehicle. Data are from one representative experiment with four animals per group (see additional data in Supporting Information, Figure S2). (B) Survival of *P. berghei*-infected mice treated as in (A). The curves represent the mean of two identical experiments, each with four animals per group (see table for statistical details). Statistical analyses were performed using unpaired Student’s t-test (parasitemia) and log-rank (Mantel-Cox) test (survival); *p< 0.05. If not indicated, no significant differences were observed.

### 3.4 Enhancement of Atovaquone and Proguanil activity by other 4-Hydroxybenzoate Analogs

The previous results spurred our interest in finding molecule elements that could be responsible for the effects on AV or Proguanil activity observed. We tackled this by using other 4-HB analogs. The first step was to create a collection of compounds to be explored (Figure 4A). This collection included 4-HB analogs with modifications to the radical bound to C4, since these have previously shown inhibitory activity in UQ biosynthesis in other organisms (Pierrel, 2017): 4-NB, 4-Bromobenzoate (4-Chlorobenzoate), carzenide (4-Sulfamoylbenzoic acid), and p-aminobenzoic acid (pABA). Additionally, we included another analog with similar characteristics, 4-Bromobenzoate (4-BrB) or with radicals introduced in other positions of the phenyl ring: methyl 4-hydroxybenzoate, para-aminosalicylic acid (PAS), 2-methyl 4-hydroxybenzoate, and 2-methyl hydroxybenzoate. Finally, as a control, we selected sulfanilamide (4-aminobenzenesulfonamide), a structurally related compound to 4-NB with no reported inhibitory activity against 4-HPT enzymes. The IC_50_ results at 48 hours (Figure 4B) revealed low antiplasmodial activity for all compounds, ranging from 1–10 mM. For sulfanilamide, the observed antiplasmodial activity was extremely low, and the R^2^ values did not indicate a clear dose-response relationship. Slightly higher antiplasmodial activity was observed for 4-HB analogs with substitutions to the radical in the C4 position. Within this group, ranked from least to most effective, the compounds classified as follows: carzenide (IC_50_ = 1.913 ± 0.01 mM), followed by 4-NB (IC_50_ = 2.55 ± 0.14 mM), 4-ClB (IC_50_ = 2.88 ± 0.92 mM), and 4-BrB (IC_50_= 9.21 ± 1.57 mM). A review of the literature revealed that two compounds in the collection had been previously tested in the parasite: PAS (Zhang et al., 1992) and sulfanilamide (Coggeshall, 1938). PAS has been described as a potent inhibitor of exogenous pABA transport (IC_50_ for transport of approximately 200 nM).

**Figure 4.**
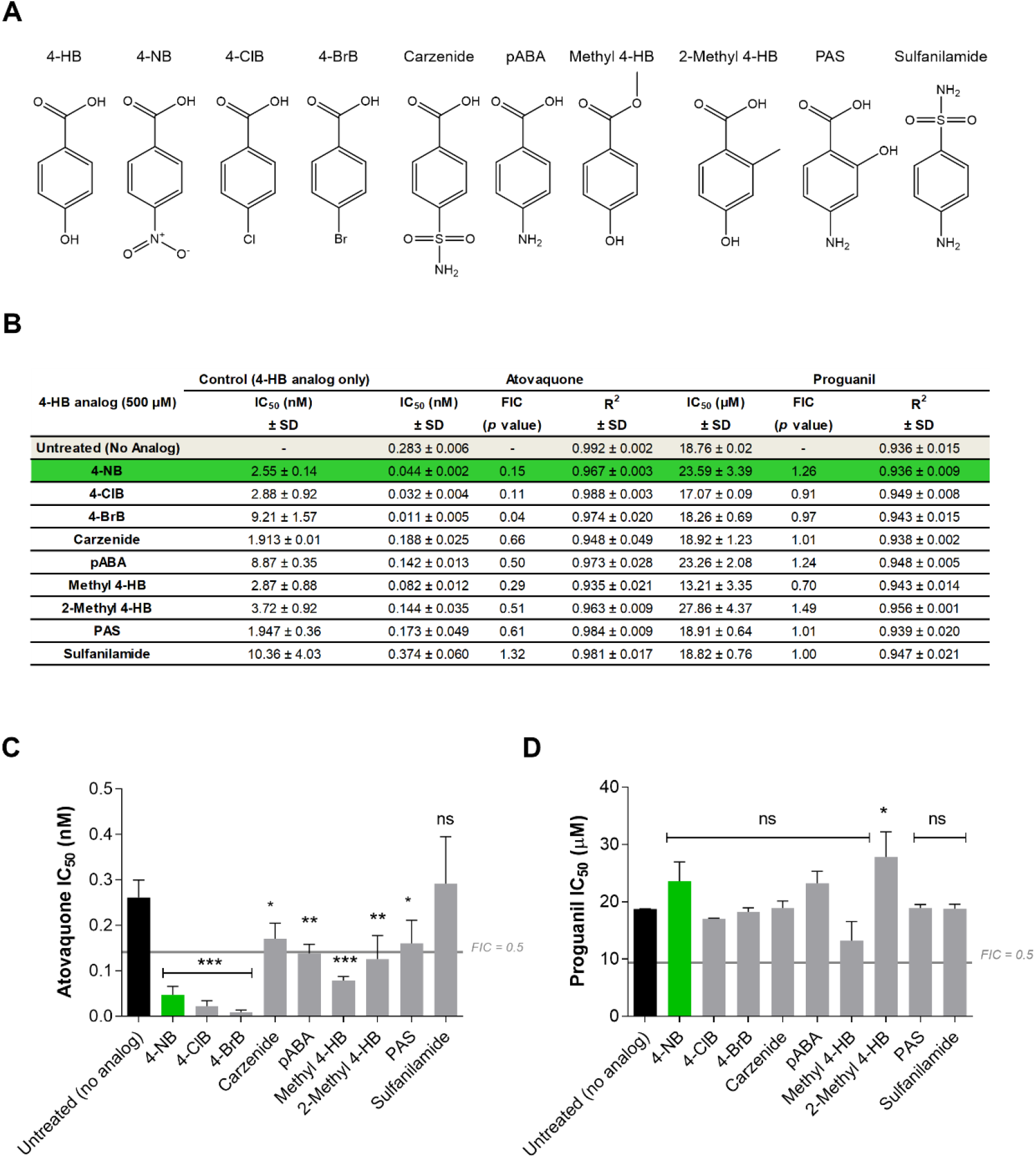
4-Hydroxybenzoate analogs effects on atovaquone and proguanil activity. (A) Structures of each of the 4-HB analog tested. (B) Values of the IC_50_ ± SD and R2 values for each 4-HB analog, as well as IC_50_ values of AV and Proguanil in routine medium or medium supplemented with each compound, as indicated. Results represent the mean of three independent experiments. Green highlights 4-NB. (C & D) Panels C and D display the results shown in Table B for AV and Proguanil, respectively. Results were analyzed using one-way analysis of variance (ANOVA) with Dunnett’s multiple comparison test compared to the control. *, P< 0.05; **, P< 0.01; ***, P< 0.001.

We then conducted pharmacological combination assays with Proguanil and AV. To this end, the IC_50_ values for Proguanil, and AV in the presence of 0.5 mM of each 4-HB analog (Figure 4B-D) were calculated. A concentration of 0.5 mM was chosen because none of the compounds in our collection caused any effect on parasite proliferation after 48 hours at that concentration. We observed pharmacological potentiation phenomena (individual FIC < 0.5) of AV using methyl 4-HB, 2-methyl 4-HB, and all 4-HB analogs with modifications to the radical in the C4 position, except for carzenide. The compounds that showed the greatest potentiation effects on AV activity were 4-NB, 4-ClB, and 4-BrB (FICs ranging from 0.05 to 0.15). No potentiation phenomena were observed for any antimalarial when using sulfanilamide (control), and only a reduction of approximately 30% in the IC_50_ value of Proguanil was achieved with methyl 4-HB We also observed that, overall, the most significant potentiation effects on AV activity were achieved by 4-HB analogs with simple modifications (-Br,-Cl,-NO_2_) to the radical in the C4 position. Additionally, methyl 4-HB and 2-methyl 4-HB also showed some potential to enhance the effects of the AV.

## 4. Discussion

AV is a current treatment for malaria. Its mechanism relies on competing with UQH_2_ for binding at cytochrome bc1 (Vaidya et al., 2000). Its toxic effects on the parasite are thus due to inhibition of UQ redox-regeneration, a process required for the activity of the DHODH enzyme involved in pyrimidine biosynthesis. However, parasites have developed resistance to AV and its combination therapy with proguanil (Vaidya et al., 2000; Färnert et al., 2003). Our group previously found that malaria parasites can be protected from AV toxicity by synthetic UQ analogs, suggesting that, conversely, decreasing UQ biosynthesis could improve AV efficacy (Verdaguer et al., 2021). Exploring this possibility, we identified 4-NB as an inhibitor of UQ biosynthesis with poor antimalarial potency on its own (IC_50_ > 2 mM at 48 h) but which strongly potentiated AV activity *in vitro* (Verdaguer et al., 2021). In this article, we questioned whether the molecular target of 4-NB could be the 4-HPT and if its AV-potentiation effect could be exploitable for *in vivo* use.

Indeed, 4-NB inhibited the 4-HPT enzymatic activity in mutant strains of *S. cerevisiae* complemented with *Pf*COQ2 in a competitive manner with the aromatic substrate. It should be noted, however, that while 4-NB can inhibit UQ biosynthesis and 4-HPT in malaria parasites, it should not be considered a specific tool for studying the effects of UQ depletion. Previous work showed that 4-NB may also affect other mitochondrial processes (Verdaguer et al., 2021). Even so, 4-NB effects on UQ metabolism seem to be a rational explanation for the observed potentiation of AV while preserving proguanil-AV synergy. Importantly, this preservation of the synergistic effect strongly suggests that the mechanism through which proguanil potentiates AV – despite still not being well understood – is different from the mechanism by which 4-NB achieves the same effect. Besides *Plasmodium* parasites, 4-NB was shown to interfere with UQ biosynthesis in highly diverse organisms, such as animal cells, cyanobacteria, and bacteria (Forsman et al., 2010; Quinzii et al., 2012; Nowicka et al., 2016; Pierrel, 2017). In human fibroblasts, the compound efficiently inhibited UQ biosynthesis, but its cytotoxicity only became evident after incubating the cells for several days at concentrations around 3–4 mM (Quinzii et al., 2012). This low toxicity of the compound is not surprising, considering that several cell types can sustain ATP production through glycolysis. Furthermore, in certain animal and plant tissues, UQ has a prolonged half-life of up to 100 hours (Wanke et al., 2000; Thelin et al., 1992). In *Plasmodium*, the effects of 4-NB are also time-dependent (Verdaguer et al., 2021). At any rate, 4-NB showed the ability to inhibit mammalian 4-HPT enzymes and UQ biosynthesis (Forsman et al., 2010). All this led us to evaluate its toxicity in combination with AV in our context. However, the presence of 0.5 mM 4-NB did not significantly reduce the CC_50_ value of AV in the LLC-MK2 cell line, reinforcing the interpretation that 4-NB could be used to potentiate AV antimalarial selectivity against animal cells. Furthermore, our experiments using AV plus 4-NB in combination in a murine malaria model were consistent with the *in vitro* results and 4-NB improved the effects of administering suboptimal doses of AV in mice. At the same time, 4-NB *per se* did not show any antimalarial effects but was well tolerated by animals. In addition to the potential combinatorial use of 4-NB and AV, these findings highlighted the importance of UQ biosynthesis for malaria parasites and suggested that PfCOQ2 could be a therapeutic target to enhance the efficacy of AV. In that respect, 4-NB could be a starting point for designing molecules capable of boosting the efficacy of AV or AV plus proguanil combinations for malaria treatment.

As mentioned, in this study, we focused on 4-NB because it had been previously tested in *Plasmodium* and has become the reference tool for studying the effects of UQ depletion. Indeed, 4-NB has already demonstrated its ability to interfere with UQ biosynthesis in highly diverse organisms such as animal cells, cyanobacteria, and bacteria (Forsman et al., 2010; Quinzii et al., 2012; Nowicka et al., 2016; Pierrel, 2017). In human fibroblasts, the compound efficiently inhibited UQ biosynthesis, but its cytotoxicity only became evident after incubating the cells for several days at concentrations around 3–4 mM (Quinzii et al., 2012). This low toxicity of the compound is not surprising, considering that several cell types can sustain ATP production through glycolysis. Furthermore, in certain animal and plant tissues, UQ has a prolonged half-life of up to 100 hours (Wanke et al., 2000; Thelin et al., 1992). In *Plasmodium*, the effects of 4-NB are also time-dependent (Verdaguer et al., 2021).

Beyond 4-NB, other authors have tested similar compounds as tools to inhibit 4-HPT activity in other organisms (Pierrel, 2017). Evidence suggesting that 4-HB analogs act on 4-HPT activity in some model organisms includes: (i) inhibitory effects on UQ formation and (ii) reversal of toxic effects and/or UQ biosynthesis inhibition by the addition of exogenous 4-HB, UQ, or dUQ (Quinzii et al., 2012; Pierrel, 2017). Only Alam et al. in 1975 and Nowicka et al. in 2016 demonstrated that 4-HB analogs directly affect 4-HPT enzymatic activity in mitochondrial preparations from animal tissues and cyanobacteria, respectively. Nowicka et al. only tested 4-NB, while Alam et al. tested acetyl salicylate, 5-methyl salicylate, 5-methoxy salicylate, 4-chlorobenzoate (4-ClB), pABA, 4-sulfamoylbenzoate (carzenide), 4-hydroxymercuribenzoate, procaine (2-diethylaminoethyl 4-aminobenzoate), and some catecholamines identified as potential regulators of 4-HPT activity: serotonin, dopamine, and norepinephrine. Nowicka et al. showed that 4-NB inhibited the formation of prenylated products from [^14^C] 4-HB, while Alam et al. demonstrated this inhibition using compounds with modifications in the C4 radical of 4-HB. Among these compounds, pABA also reduced the formation of prenylated products from [^14^C] 4-HB, which led Alam et al. to further investigate this phenomenon. The authors demonstrated that, similar to 4-HB, pABA can also be prenylated by polyprenyltransferases in animal tissues. However, the resulting product, 3-polyprenyl-pABA, cannot be deaminated to continue in UQ biosynthesis. In contrast, yeasts can form UQs from pABA because, in addition to prenylating pABA, they can also deaminate 3-polyprenyl-pABA (Marbois et al., 2010).

Considering the above, the literature concludes that, for the 4-HPT enzyme to prenylate a given aromatic molecule, the substrate must meet two requirements: the presence of a radical in the C4 position with the ability to transfer electrons (-OH, -NH_2_, among others) and a strong electron-attracting group in the C1 position (the carboxyl group) to generate sufficient electron density at the C3 position (Pierrel, 2017). This electron density allows for a nucleophilic attack on the oxygen-carbon bond of the polyprenyl pyrophosphate. However, the size and electronegativity of radicals in the C4 position may also reduce nucleophilic interaction at the C3 position, making alkylation unfeasible. In line with this reasoning, this study observed pharmacological potentiation phenomena of AV using methyl 4-HB, 2-methyl 4-HB, and all 4-HB analogs with modifications in the C4 radical, except for carzenide. The compounds showing the most significant potentiation effects on AV activity were 4-NB, 4-ClB, and 4-BrB, which remarked the importance of an electron-drawing residue at C4. No potentiation phenomena were observed for any antimalarial when using sulfanilamide (control), and only a reduction of approximately 30% in Proguanil’s IC_50_ value was achieved with methyl 4-HB. These results further suggest that the mitochondrial target of Proguanil is likely not directly related to UQ metabolism. We also concluded that, overall, the most significant potentiation effects on AV activity are achieved by 4-HB analogs with simple modifications (-Br, -Cl, -NO_2_) to the radical in the C4 position. Additionally, methyl 4-HB and 2-methyl 4-HB also showed some potential to enhance AV’s effects. A review of the literature revealed that two compounds in the collection had been previously tested in parasites: PAS (Zhang et al., 1992) and sulfanilamide (Coggeshall, 1938). No evidence for a UQ-related mechanism has been reported for any of them. Sulfanilamide is a known folate biosynthesis inhibitor, while PAS has been described as a potent inhibitor of exogenous pABA transport (IC_50_ for transport of approximately 200 nM) (Zhang et al., 1992). In agreement with this, we found no potentiation effect of those two compounds on AV action. Interestingly, a slight potentiation effect on AV activity was also observed using pABA. It is important to note that pABA is a metabolite produced by the parasite and is present in human blood (Ansbacher, 1941). Taylor in 1957 showed that supplementing the diet of chicks infected with *P. gallinaceum* with high doses of pABA reduced parasitemia levels. However, this antimalarial effect of pABA could be reversed by adding 4-HB to the diet, indicating that pABA may interrupt UQ biosynthesis.

Finally, it should be noticed that AV and similar analogs are already used in the treatment of other diseases, such as babesiosis (Pudney et al., 1997) and toxoplasmosis (Montazeri et al., 2018), and are being investigated for the treatment of leishmaniasis (Croft et al., 1992), among other infectious diseases. Therefore, the conclusions and strategy used in this work to enhance AV efficacy may also be of interest to many other diseases.

## 5. Conclusions

Our study showed that 4-NB effectively inhibited 4-HPT in *Plasmodium falciparum*, this being the likely reason for the previously observed decrease UQ biosynthesis in malaria parasites. This inhibition not only enhanced the antiplasmodial efficacy of AV but also increased its selectivity compared to animal cells, without detrimental effects on proguanil efficacy. Furthermore, the combination of 4-NB and AV significantly improved therapeutic outcomes in a murine malaria model, remarking the potential of 4-HB analogs as an adjuvant to current antimalarial therapies. Finally, testing various 4-HB analogs allowed us to confirm some of the chemical requirements for these compounds to potentiate AV activity. These findings underscored the importance of targeting UQ biosynthesis in malaria parasites and may provide a promising strategy to overcome resistance and enhance the efficacy of existing antimalarial drugs.

## ACKNOWLEDGMENTS

The authors would like to thank Sírio Libanês Hospital (NESTA, São Paulo, Brazil) for the gift of erythrocytes. The authors also thank Dr. M. H. de Barros (Dept. Microbiology, University of São Paulo) for the gift of *ΔCOQ2* strain of *S. cerevisiae* and its parental W303-1a.

## AUTHORS CONTRIBUTION

IBV, MC, GOC, MFS, MAP, MMF contributed to conceptualization, formal analysis, investigation, methodology and writing. AMK, AHL, MC also contributed in project administration, funding acquisition, supervision and writing – review & editing.

## ETHICS STATEMENT

This study was carried out in strict accordance with the recommendations provided by the Guide for the Care and Use of Laboratory Animals of the Brazilian National Council of Animal Experimentation (http://www.cobea.org.br/). The protocol was approved by the Committee on the Ethics of Animal Experiments of the Institutional Animal Care and Use Committee at the Institute of Biomedical Sciences of the University of São Paulo (Protocol nº 2420280823).

## REFERENCES

1. Alam SS, Nambudiri AM, Rudney H. J-Hydroxybenzoate: polyprenyl transferase and the prenylation of 4-aminobenzoate in mammalian tissues. Arch Biochem Biophys. 1975;171(1):183–90.

2. Ansbacher S. p-aminobenzoic acid, a vitamin. Science. 1941;93(2407):164–5.

3. Arodin Selenius, L., Wallenberg Lundgren, M., Jawad, R., Danielsson, O., & Björnstedt, M. (2019). The cell culture medium affects growth, phenotype expression and the response to selenium cytotoxicity in A549 and HepG2 cells. Antioxidants, 8(5), 130.

4. Coggeshall LT. The cure of Plasmodium knowlesi malaria in rhesus monkeys with sulfanilamide and their susceptibility to reinfection. The American Journal of Tropical Medicine and Hygiene 1938;s1-18(6):715–21.

5. Crispim, M., Verdaguer, I. B., Silva, S. F., & Katzin, A. M. (2022). Suitability of methods for Plasmodium falciparum cultivation in atmospheric air. Memorias do Instituto Oswaldo Cruz, 117, e210331.

6. Croft, S. L., Hogg, J., Gutteridge, W. E., Hudson, A. T., & Randall, A. W. (1992). The activity of hydroxynaphthoquinones against Leishmania donovani. Journal of Antimicrobial Chemotherapy, 30(6), 827–832.

7. de Macedo, C. S., Uhrig, M. L., Kimura, E. A., & Katzin, A. M. (2002). Characterization of the isoprenoid chain of coenzyme Q in Plasmodium falciparum. FEMS microbiology letters, 207(1), 13–20.

8. Desjardins, R. E., Canfield, C. J., Haynes, J. D., & Chulay, J. D. (1979). Quantitative assessment of antimalarial activity in vitro by a semiautomated microdilution technique. Antimicrobial agents and chemotherapy, 16(6), 710–718.

9. Dunn, BarbaraB.,, and C. Richard Wobbe. “Preparation of protein extracts from yeast.” Current protocols in molecular biology 23.1 (1993): 13–13.

10. Ellis, J. E. (1994). Coenzyme Q homologs in parasitic protozoa as targets for chemotherapeutic attack. Parasitology Today, 10(8), 296–301.

11. Färnert, Anna, et al. “Evidence of Plasmodium falciparum malaria resistant to AV and proguanil hydrochloride.” Bmj 326.7390 (2003): 628–629.

12. Farooq, U., & Mahajan, R. C. (2004). Drug resistance in malaria. Journal of vector borne diseases, 41(3/4), 45.

13. Fiorillo M, Lamb R, Tanowitz HB, Mutti L, Krstic-Demonacos M, Cappello AR, et al. Repurposing atovaquone: targeting mitochondrial complex III and OXPHOS to eradicate cancer stem cells. Oncotarget 2016;7(23):34084–99.

14. Forsman, Ulrika, et al. “4-Nitrobenzoate inhibits coenzyme Q biosynthesis in mammalian cell cultures.” Nature chemical biology 6.7 (2010): 515–517.

15. Gietz, R. D., & Woods, R. A. (2002). Transformation of yeast by lithium acetate/single-stranded carrier DNA/polyethylene glycol method. In Methods in enzymology (Vol. 350, pp. 87–96). Academic Press.

16. Kissin, I. (2013). An early indicator of drug success: Top Journal Selectivity Index. Drug Design, Development and Therapy, 93–98.

17. Lambros, C., & Vanderberg, J. P. (1979). Synchronization of Plasmodium falciparum erythrocytic stages in culture. The Journal of parasitology, 418–420.

18. Mandel, P., and P. Metais. “Comptes rendus des seances de la Societe de biologie et de ses filiales.” Journal de la Société de Biologie 142 (1948).

19. Marbois B, Xie LX, Choi S, Hirano K, Hyman K, Clarke CF. para-Aminobenzoic acid is a precursor in coenzyme Q6 biosynthesis in Saccharomyces cerevisiae. J Biol Chem 2010;285(36):27827–38.

20. McKeage, Kate, and Lesley J. Scott. “AV/proguanil.” Drugs 63.6 (2003): 597–623.

21. Melzer, Martin, and Lutz Heide. “Characterization of polyprenyldiphosphate: 4-hydroxybenzoate polyprenyltransferase from Escherichia coli.” Biochimica et Biophysica Acta (BBA)-Lipids and Lipid Metabolism 1212.1 (1994): 93–102.

22. Montazeri, M., Mehrzadi, S., Sharif, M., Sarvi, S., Shahdin, S., & Daryani, A. (2018). Activities of anti-Toxoplasma drugs and compounds against tissue cysts in the last three decades (1987 to 2017), a systematic review. Parasitology research, 117, 3045–3057.

23. Mosmann, T. (1983). Rapid colorimetric assay for cellular growth and survival: application to proliferation and cytotoxicity assays. Journal of immunological methods, 65(1-2), 55–63.

24. Nikfarjam, L., & Farzaneh, P. (2012). Prevention and detection of Mycoplasma contamination in cell culture. Cell Journal (Yakhteh), 13(4), 203.

25. Nowicka B, Kruk J. Cyanobacteria use both p-hydroxybenozate and homogentisate as a precursor of plastoquinone head group. Acta Physiol Plant 2016;38(2):49.

26. Nuralitha, S., Murdiyarso, L. S., Siregar, J. E., Syafruddin, D., Roelands, J., Verhoef, J., … & Marzuki, S. (2017). Within-host selection of drug resistance in a mouse model reveals dose-dependent selection of atovaquone resistance mutations. Antimicrobial Agents and Chemotherapy, 61(5), 10–1128.

27. Painter, H. J., Morrisey, J. M., Mather, M. W., & Vaidya, A. B. (2007). Specific role of mitochondrial electron transport in blood-stage Plasmodium falciparum. nature, 446(7131), 88–91.

28. Pfaff, C., Glindemann, N., Gruber, J., Frentzen, M., & Sadre, R. (2014). Chorismate pyruvate-lyase and 4-hydroxy-3-solanesylbenzoate decarboxylase are required for plastoquinone biosynthesis in the cyanobacterium Synechocystis sp. PCC6803. Journal of Biological Chemistry, 289(5), 2675–2686.

29. Pierrel, Fabien. “Impact of chemical analogs of 4-hydroxybenzoic acid on coenzyme Q biosynthesis: from inhibition to bypass of coenzyme Q deficiency.” Frontiers in Physiology 8 (2017): 436.

30. Quinzii, Catarina M., et al. “Effects of inhibiting CoQ10 biosynthesis with 4-nitrobenzoate in human fibroblasts.” PloS one 7.2 (2012): e30606.

31. Radloff, Paul D., et al. “AV and proguanil for Plasmodium falciparum malaria.” The Lancet 347.9014 (1996): 1511–1514.

32. Rowe, J. A., Scragg, I. G., Kwiatkowski, D., Ferguson, D. J., Carucci, D. J., & Newbold, C. I. (1998). Implications of mycoplasma contamination in Plasmodium falciparum cultures and methods for its detection and eradication. Molecular and biochemical parasitology, 92(1), 177–180.

33. Sies, H., & Packer, L. (2004). Quinones and Quinone Enzymes, Part B. Elsevier.

34. Smilkstein, M., Sriwilaijaroen, N., Kelly, J. X., Wilairat, P., & Riscoe, M. (2004). Simple and inexpensive fluorescence-based technique for high-throughput antimalarial drug screening. Antimicrobial agents and chemotherapy, 48(5), 1803–1806.

35. Srivastava, I. K., & Vaidya, A. B. (1999). A mechanism for the synergistic antimalarial action of AV and proguanil. Antimicrobial agents and chemotherapy, 43(6), 1334–1339.

36. Taylor, A. E. (1957). The effect of paraminobenzoic acid, parahydroxybenzoic acid and riboflavin on Plasmodium gallinaceum in chicks. Transactions of the Royal Society of Tropical Medicine and Hygiene, 51, 241–247.

37. Thelin A, Schedin S, Dallner G. Half-life of ubiquinone-9 in rat tissues. FEBS Lett 1992;313(2):118–20.

38. Trager, W., & Jensen, J. B. (1976). Human malaria parasites in continuous culture. Science (New York, N.Y.), 193(4254), 673–675.

39. Vaidya, Akhil B., and Michael W. Mather. “AV resistance in malaria parasites.” Drug Resistance Updates 3.5 (2000): 283–287.

40. Valenciano, A. L., Fernández-Murga, M. L., Merino, E. F., Holderman, N. R., Butschek, G. J., Shaffer, K. J., … & Cassera, M. B. (2019). Metabolic dependency of chorismate in Plasmodium falciparum suggests an alternative source for the ubiquinone biosynthesis precursor. Scientific reports, 9(1), 1–13.

41. Van Dooren, Giel G., Luciana M. Stimmler, and Geoffrey I. McFadden. “Metabolic maps and functions of the Plasmodium mitochondrion.” FEMS microbiology reviews 30.4 (2006): 596–630.

42. Verdaguer, I. B., Crispim, M., Zafra, C. A., Sussmann, R. A. C., Buriticá, N. L., Melo, H. R., … & Katzin, A. M. (2021). Exploring ubiquinone biosynthesis inhibition as a strategy for improving atovaquone efficacy in malaria. Antimicrobial Agents and Chemotherapy, 65(4), 10–1128.

43. Verdaguer, I. B., Zafra, C. A., Crispim, M., Sussmann, R. A., Kimura, E. A., & Katzin, A. M. (2019). Prenylquinones in human parasitic protozoa: biosynthesis, physiological functions, and potential as chemotherapeutic targets. Molecules, 24(20), 3721.

44. Wanke M, Swiezewska E, Dallner G. Half-life of ubiquinone and plastoquinone in spinach cells. Plant Sci 2000;154(2):183–7.

45. World Health Organization, World malaria report 2023, WHO. (2023).

46. Zafra, C. A., Crispim, M., Verdaguer, I. B., Ríos, A. G., Moura, G. C., Katzin, A. M., & Hernández, A. (2023). Plasmodium falciparum COQ2 gene encodes a functional 4-hydroxybenzoate polyprenyltransferase. FEMS Microbiology Letters, fnad050.

47. Zhang Y, Merali S, Meshnick SR. p-Aminobenzoic acid transport by normal and Plasmodium falciparum-infected erythrocytes. Mol Biochem Parasitol 1992;52(2):185– 94.

