## Supporting Information Figures for "4-Nitrobenzoate inhibits 4-hydroxybenzoate polyprenyltransferase in malaria parasites and enhances atovaquone efficacy"

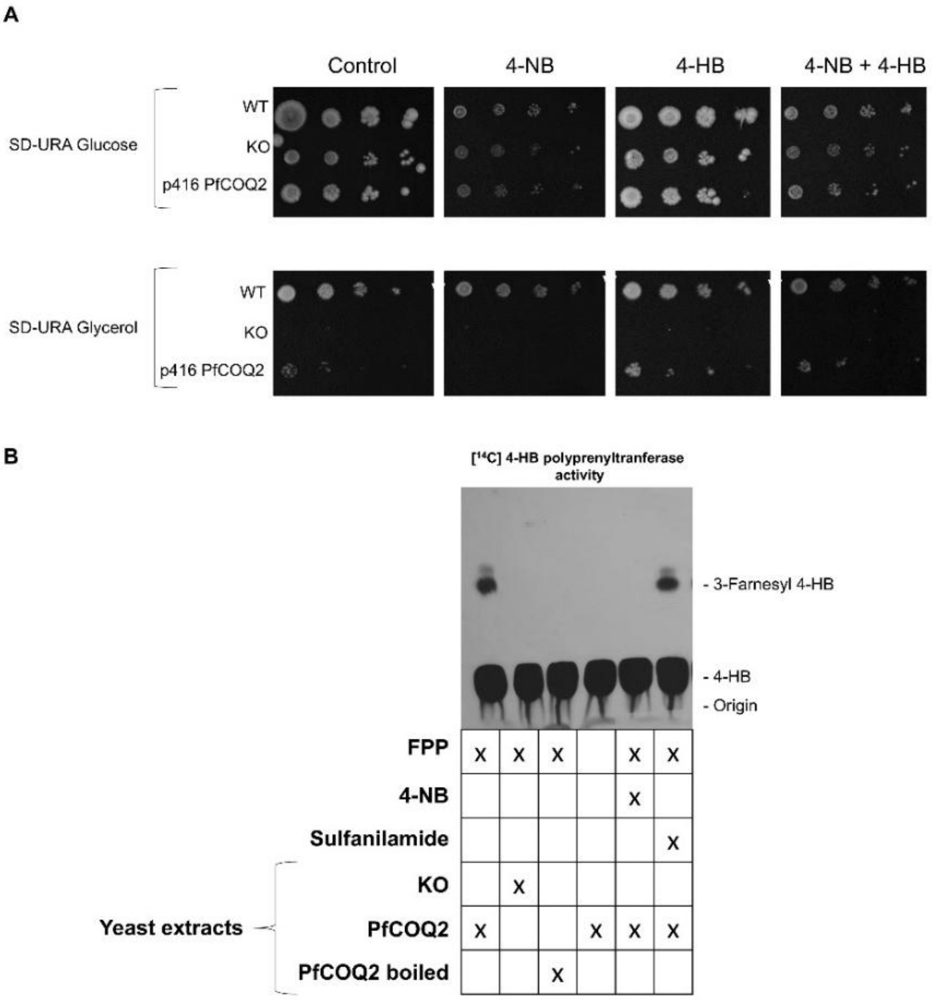


**Figure S1.** ***PfCOQ2* complementation and enzymatic activity.** (A) The figure shows the growth of yeast strains in SD + glucose / SD + glycerol media containing a concentration of 1 mM of different drugs, as indicated. This experiment was performed three times with similar results. (B) The figure shows the analysis of 4-HPT enzymatic activity in *Pf*COQ2 complemented yeasts under the presence of 0.5 mM of different drugs. The compounds added to the enzymatic reaction are indicated on the TLC autoradiography. The retention of different compounds is also indicated. This experiment was performed three times with similar results.

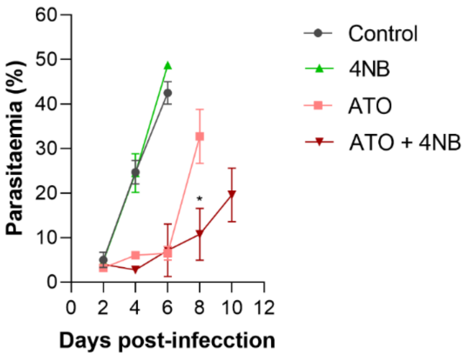


**Figure S2. 4-Nitrobenzoate improves the efficacy of atovaquone in vivo.** Parasitemia in *P. berghei* ANKA-infected mice treated with atovaquone (AV), 4-nitrobenzoate (4-NB), or their combination. All groups received the same amount of vehicle. Data are from one representative experiment with four animals per group. Statistical analyses were performed using unpaired Student's t-test (parasitemia) and log-rank (Mantel-Cox) test (survival); *p<0.05. If not indicated, no significant differences were observed
